# Convergent evolution of effector protease recognition by Arabidopsis and barley

**DOI:** 10.1101/374264

**Authors:** Morgan E. Carter, Matthew Helm, Antony Chapman, Emily Wan, Ana Maria Restrepo Sierra, Roger W. Innes, Adam J. Bogdanove, Roger P. Wise

## Abstract

The *Pseudomonas syringae* cysteine protease AvrPphB activates the Arabidopsis resistance protein RPS5 by cleaving a second host protein, PBS1. AvrPphB induces defense responses in other plant species, but the genes and mechanisms mediating AvrPphB recognition in those species have not been defined. Here, we show that AvrPphB induces defense responses in diverse barley cultivars. We show also that barley contains two *PBS1* orthologs, that their products are cleaved by AvrPphB, and that the barley AvrPphB response maps to a single locus containing a nucleotide-binding leucine-rich repeat (NLR) gene, which we termed *Avr<u>P</u>ph<u>B</u> Resistance <u>1</u>* (*Pbr1*). Transient co-expression of PBR1 with wild-type AvrPphB, but not a protease inactive mutant, triggered defense responses, indicating that PBR1 detects AvrPphB protease activity. Additionally, PBR1 co-immunoprecipitated with barley and *N. benthamiana* PBS1 proteins, suggesting mechanistic similarity to detection by RPS5. Lastly, we determined that wheat cultivars also recognize AvrPphB protease activity and contain a *Pbr1* ortholog. Phylogenetic analyses showed however that *Pbr1* is not orthologous to *RPS5*. Our results indicate that the ability to recognize AvrPphB evolved convergently, and imply that selection to guard PBS1-like proteins is ancient. Also, the results suggest that PBS1-based decoys may be used to engineer protease effector recognition-based resistance in barley and wheat.

## Introduction

Plant disease resistance is often mediated by intracellular innate immune receptors known as nucleotide-binding leucine-rich repeat proteins (NLRs). The primary function of NLRs is to detect the presence of pathogen-secreted effector proteins, sometimes indirectly through effector-induced modification of other host proteins (Jones and Dangl, 2006). Recognition of effectors by NLRs usually activates a programmed cell death response known as the hypersensitive reaction (HR) (Z Klement and Goodman, 1967; Coll et al., 2011). A well-studied example of an NLR that indirectly detects its cognate effector is RPS5 from Arabidopsis. RPS5 detects the *Pseudomonas syringae* pv. *phaseolicola* effector protease AvrPphB by monitoring the conformational status of an Arabidopsis substrate of AvrPphB, the serine/threonine protein kinase PBS1 (Shao et al., 2003; Ade et al., 2007; DeYoung et al., 2012; Qi et al., 2014). RPS5 forms a “pre-activation complex” with PBS1, and when PBS1 is cleaved by AvrPphB, the resulting conformational change is sensed by RPS5, culminating in activation of the NLR and subsequent induction of HR (Shao et al., 2003; Ade et al., 2007; DeYoung et al., 2012).

*PBS1* is one of the most well conserved defense-related genes in flowering plants, and the products of *PBS1* orthologs in wheat and Arabidopsis can be cleaved by AvrPphB (Caldwell and Michelmore, 2009; Kim et al., 2016; Sun et al., 2017). PBS1 belongs to receptor-like cytoplasmic kinase (RLCK) family VII, which has many members with demonstrated roles in pattern-triggered immunity (PTI) (Zhang et al., 2010; DeYoung et al., 2012). For example, family VII RLCKs BIK1 and PBL1 physically associate with the flagellin-detecting receptor FLS2 (Zhang et al., 2010).. Of the 45 Arabidopsis proteins within RLCK family VII, 9 are PBS1-like (PBL) kinases cleaved by AvrPphB (Shao et al., 2003; Zhang et al., 2010; DeYoung et al., 2012), and AvrPphB indeed inhibits FLS2-dependent PTI, as well as defense responses triggered by EfTu and chitin (Zhang et al., 2010). However, only its cleavage of PBS1 activates RPS5 (Ade et al., 2007; Zhang et al., 2010).

Because of their role in PTI, RLCKs and other kinases are commonly targeted by pathogen effectors (Yamaguchi et al., 2013). Examples beyond the AvrPphB-PBS1 interaction include the RLCK BIK1, which is uridylylated by AvrAC from *Xanthomonas campestris* pv. *campestris*, and the receptor-like kinase BAK1 which is bound by the *Pseudomonas syringae* pv. *tomato* effectors AvrPto and AvrPtoB to inhibit signaling (Shan et al., 2008; Feng et al., 2012). Some kinases targeted by effectors appear to play little to no primary role in immunity, but function as decoys, guarded by NLRs to detect effector activity. An example is the RLCK PBL2 in Arabidopsis, which is uridylylated by AvrAC like BIK1, and is guarded by the NLR ZAR1 (Wang et al., 2015).

Determining whether PBS1 orthologs are guarded in diverse plant species is of particular interest because it will provide insight into the evolution of disease resistance gene specificity and could enable engineering of new disease resistance specificities in crop plants. Kim et al. (2016) demonstrated that the AvrPphB cleavage site sequence within PBS1 can be substituted with a sequence recognized by an effector protease of another pathogen, thereby generating a synthetic PBS1 decoy. Cleavage of PBS1 decoys *in planta* activates RPS5-dependent HR, effectively broadening the recognition specificity of RPS5 (Kim et al., 2016). Thus, in plant species in which a PBS1 ortholog is guarded, engineering these orthologs to serve as substrates of other pathogen proteases offers an attractive approach for generating resistance tailored to pathogens of those species. Given that plant pathogenic viruses, bacteria, fungi, oomycetes, and nematodes express proteases during infection, engineering the RPS5/PBS1 surveillance system may be an effective strategy for developing resistance to many important plant diseases (Adams et al., 2005; Dean, 2011; Antonino de Souza Júnior et al., 2013; Jashni et al., 2015).

While it was recently reported that bread wheat (*Triticum aestivum* subsp. *aestivum*) encodes a homolog of Arabidopsis PBS1, TaPBS1, that can be cleaved by AvrPphB, it remains unknown whether TaPBS1 is guarded, *i.e*., whether wheat or other cereals can recognize and respond to AvrPphB (Sun et al., 2017). Unlike the PBL kinases, NLR genes are under intense selection pressure to diversify, and they vary greatly in number and structure across plant genomes (Jacob et al., 2013). For example, grasses are missing the entire TIR domain-containing family of NLRs that is present in many dicots (Collier et al., 2011). However, it is plausible that proteins functionally analogous to RPS5 guard AvrPphB-cleavable PBS1 homologs in the grasses, especially given the central role that RLCK proteins play in immunity. We investigated this hypothesis using diploid barley (*Hordeum vulgare* subsp. *vulgare*) as a model because of its rich genetic resources, including a high-quality genome sequence (Mascher et al., 2017) and large nested association mapping (NAM) populations (Nice et al., 2016).

Here, we show that multiple barley varieties indeed recognize and respond to AvrPphB protease activity and that barley also contains PBLs that are cleaved by AvrPphB. Using newly developed NAM resources, we mapped the AvrPphB response to a single segregating locus on chromosome 3HS and identified an NLR gene that we named *Avr<u>P</u>phB <u>R</u>esistance <u>1</u>* (*Pbr1*). We confirmed that PBR1 mediates AvrPphB recognition using transient expression assays in *Nicotiana benthamiana* and determined that PBR1 associates with PBS1 homologs *in planta*. Phylogenetic analyses indicate that *Pbr1* and *RPS5* are not orthologous, hence the ability to recognize AvrPphB protease activity has evolved independently in monocots and dicots. Lastly, we show that wheat varieties also recognize AvrPphB protease activity and harbor an ortholog of *Pbr1* in a syntenic position on chromosome 3B, suggesting that the PBS1-decoy system might be deployed in barley and in wheat.

## Results

### AvrPphB, and not a catalytically inactive derivative, triggers defense responses in barley

To test whether barley can detect AvrPphB protease activity, we delivered AvrPphB to barley leaves using *P. syringae* pathovar *tomato* strain D36E, which is a derivative of strain DC3000 lacking all type III secretion system effectors (Wei et al., 2015). Seedlings were infiltrated with D36E expressing AvrPphB or the catalytically inactive mutant AvrPphB(C98S) and scored for visible responses at 2 and 5 days post infiltration.

We tested a diverse set of barley lines and observed a variety of responses. Representative examples are shown in Fig. 1, and the complete list of cultivars and their responses are provided in Supplementary Table S1. Based on the range of responses we saw, we scored the phenotypes as no response (N) or one of 4 responses: low chlorosis (LC) indicates a weak, but noticeable response, chlorosis (C) for strong yellow, high chlorosis (HC) for a chlorotic response that gives way to cell death, and hypersensitive reaction (HR) for cell collapse and browning visible by day 2.

**Figure 1.**
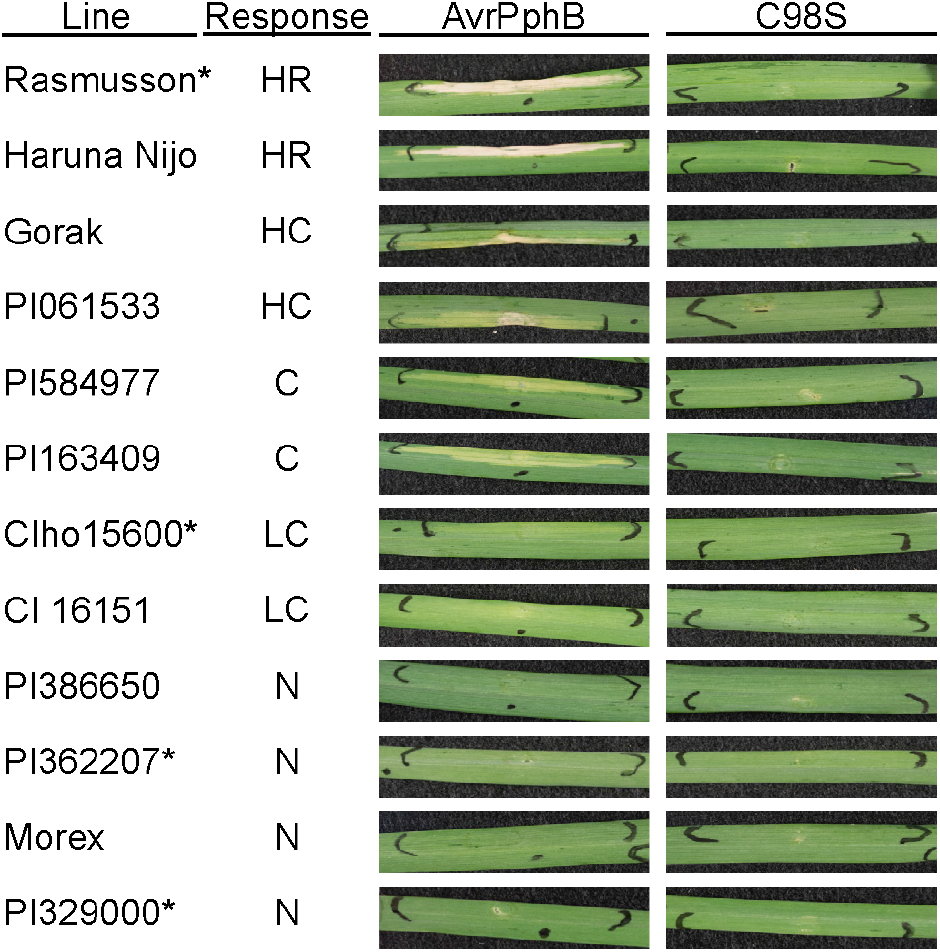
AvrPphB protease activity elicits a range of responses in barley lines. Representative barley leaves from 12 lines after infiltration with strains of *Pseudomonas syringae* pv. *tomato* DC3000(D36E) expressing AvrPphB or a catalytically inactive mutant, AvrPphB(C98S). Primary leaves of ten day old plants were infiltrated using needleless syringe with a bacterial suspension at an OD_600_=0.5 and photographed at 5 dpi. Phenotypes were scored as: N - no response; LC – low chlorosis; C – chlorosis; HC - high chlorosis; HR – hypersensitive reaction. At least six plants were infiltrated with both strains per line over two repeats. Asterisks (*) indicate parental lines of the mapping population families used for GWAS. Responses of all lines tested are recorded in Table S1.

Of the 150 barley genotypes screened, 29 were scored as LC, 17 as C, 13 as HC and 6 as HR (Supp. Table S1). Both chlorotic and cell death responses were considered defense responses, as both have been documented as such for grasses (Smith and Mansfield, 1981).

### PBS1 homologs in barley contain the AvrPphB recognition site and are cleaved by AvrPphB

Having found that many barley lines recognize D36E expressing AvrPphB, we sought to determine whether barley contains a recognition system functionally analogous to the Arabidopsis RPS5-PBS1 pathway. Because *PBS1* is one of the most well conserved defense genes in flowering plants, with orthologs present in monocot and dicot crop species (Caldwell and Michelmore, 2009), we first asked whether barley contains a PBS1 homolog cleavable by AvrPphB.

We used amino acid sequences from all characterized Arabidopsis PBS1-like (AtPBL) proteins, Arabidopsis PBS1 (AtPBS1), and twenty barley PBS1-like (HvPBL) protein sequences homologous to AtPBS1 and AtPBL proteins to identify the barley proteins most closely related to Arabidopsis PBS1. Bayesian phylogenetic analyses showed that HORVU2Hr1G070690.2 (MLOC_13277) was the closest homolog to AtPBS1, whereas HORVU3Hr1G035810.1 (MLOC_12866) was the second most closely related (Fig. 2A; Supp. Fig. S1). Both proteins are more similar to AtPBS1 than to other AtPBL and HvPBL proteins, indicating that the two barley genes are co-orthologous to *AtPBS1*. Full-length amino acid alignments showed that HORVU2Hr1G070690.2 and HORVU3Hr1G035810.1 are 66% and 64% identical to Arabidopsis PBS1, respectively (Supp. Fig. S2). Alignment of the two barley gene products and Arabidopsis PBS1 across the kinase domain showed 86% and 79% identity, respectively. Further characterization of the HvPBS1 orthologs showed that each contains several domains that are conserved in AtPBS1, including putative N-terminal palmitoylation and myristoylation sites required for plasma membrane localization and the protease cleavage site sequence recognized by AvrPphB (Fig. 2B; Supp. Fig. S2). We therefore designated HORVU2Hr1 G070690.2 (MLOC_13277) as *HvPbs1-1* and HORVU3Hr1G035810.1 (MLOC_12866) as *HvPbs1-2*.

**Figure 2.**
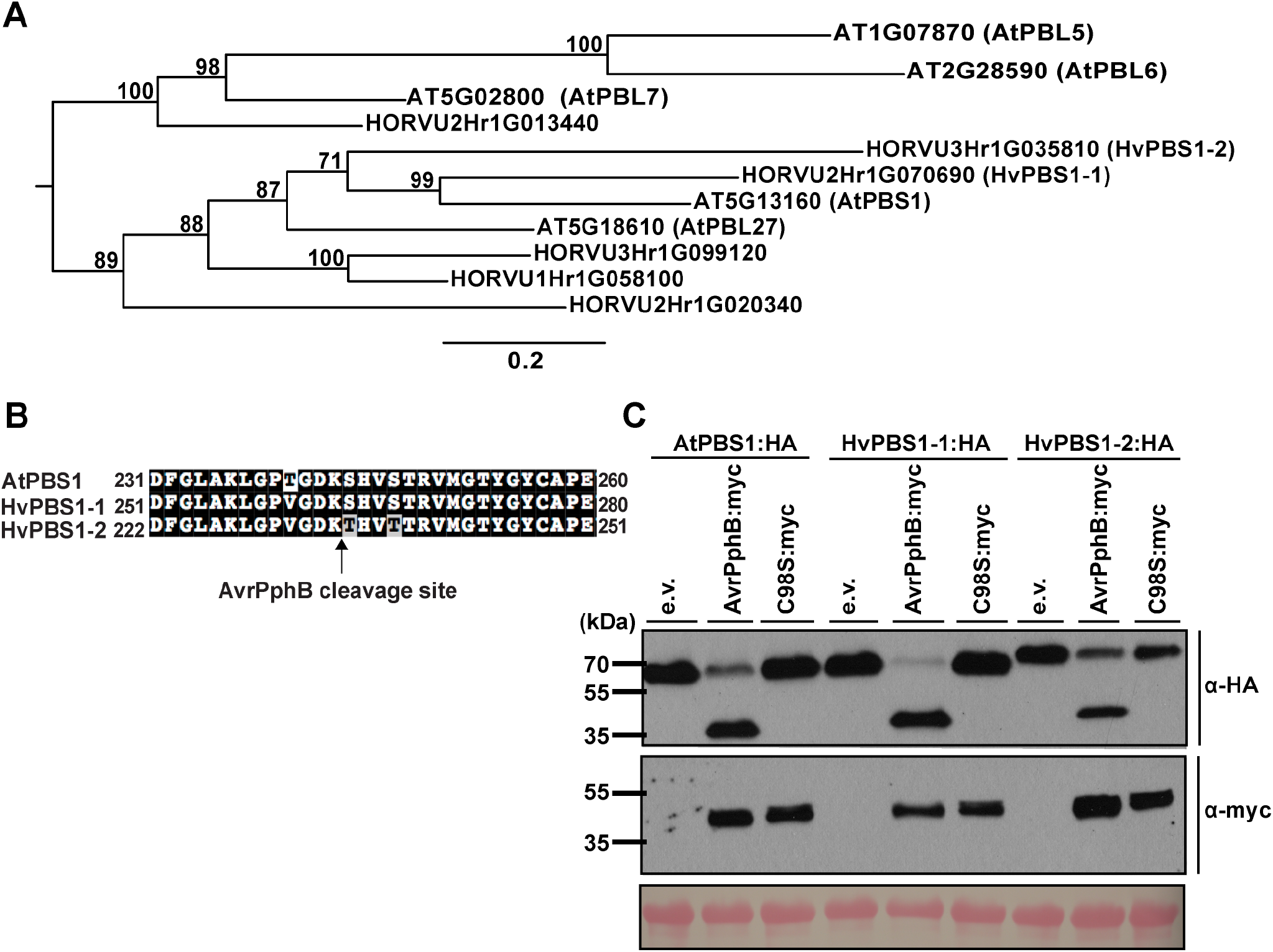
Barley contains two PBS1 homologs that are cleaved by AvrPphB. **A**) HORVU2Hr1G070690.2 (HvPBS1-1) and HORVU3Hr1G035810.1 (HvPBS1-2) are co-orthologous to Arabidopsis PBS1. Shown is a Bayesian phylogenetic tree generated from the amino acid sequences of Arabidopsis PBS1 (AtPBS1) and closely related barley homologs of AtPBS1. This tree is a subset of Supplemental Figure 1 displaying the proteins most similar to AtPBS1. Branch annotations represent Bayesian posterior probabilities as a percentage. **B**) Alignment of the activation segment sequences of AtPBS1 and the barley PBS1 homologs. The AvrPphB cleavage site is indicated by the arrow. Numbers indicate amino acid positions. **C**) Cleavage of HvPBS1-1 and HvPBS1-2 by AvrPphB. HA-tagged barley PBS1 homologs or AtPBS1 were transiently coexpressed with or without myc-tagged AvrPphB, or a protease inactive derivative [AvrPphB(C98S)] in *N. benthamiana*. Six hours post-transgene induction, total protein was extracted and immunoblotted with the indicated antibodies. Two independent experiments were performed with similar results.

Conservation of the AvrPphB cleavage site sequences within the barley PBS1 homologs led us to hypothesize that AvrPphB would cleave HvPBS1-1 and HvPBS1-2. To test this, HvPBS1-1 and HvPBS1-2 were fused to a three-copy human influenza haemagglutinin (3xHA) epitope tag and transiently co-expressed with AvrPphB:myc in *N. benthamiana*. Western blot analysis indeed showed that HvPBS1-1:HA and HvPBS1-2:HA are each cleaved by AvrPphB:myc (Fig. 2C). As a control, we co-expressed HvPBS1-1:HA and HvPBS1-2:HA with protease inactive AvrPphB(C98S):myc, and this did not produce any cleavage products (Fig. 2C). Collectively, these data show that barley contains two PBS1 homologs whose protein products can be cleaved by AvrPphB and whose function may be analogous to AtPBS1.

### A single NLR gene-rich region in the barley genome is associated with AvrPphB response

Given the response to AvrPphB in some barley lines and the presence of conserved AvrPphB-cleavable PBS1 homologs in barley, we hypothesized that the responding barley lines contain a PBS1-guarding NLR analogous to RPS5. To identify candidates, we carried out a genome wide association study (GWAS). The Rasmusson spring barley nested association mapping (NAM) population generated by the US Barley CAP (A. Ollhoff and K. Smith, University of Minnesota, unpublished) contains 6,161 RILs derived from crosses between the elite malting line Rasmusson and 88 diverse donor parents, each of which has associated SNP marker data. Rasmusson, the common parent, displays an HR when infiltrated with D36E expressing AvrPphB whereas the other parents vary in their responses (Fig. 1). For the GWAS, three NAM subpopulations (families) were chosen: two derived from non-responding parents, PI329000 (family HR656) and PI366207 (HR658), and one from a low-chlorosis response parent, CIho15600 (HR620) (Fig. 1).

As expected for a qualitative, single gene trait, the responses segregated ~1:1 within each family of RILs; 39 of 73 HR656 lines, 19 of 36 HR658 lines, and 29 of 66 HR620 lines displayed an HR following infiltration with D36E expressing AvrPphB (a total of 87 out of 175 RILs tested; Supp. Table S2). Co-segregation of AvrPphB response with SNPs was analyzed using the R/NAM package (Xavier et al., 2015), which included 13,981 SNPs in the analysis of the 175 lines (see Methods). The GWAS identified a 22.65 Mb region between positions 660,376,398 (SNP 3H2_266065765) and 683,030,529 (SNP 3H2_288719896) on the short arm of chromosome 3H associated with AvrPphB response (Fig. 3A). Neither *HvPbs1-1* nor *HvPbs1-2* are in this region, supporting the hypothesis that an NLR, and not a PBS1 homolog, is the determinant of AvrPphB response. Notably, analysis of each of the three families individually identified the same locus on chromosome 3H as the only significant association (Fig. 3B). The most significant SNP and the number of SNPs used in the analysis varied by population due to the SNP variation between Rasmusson and each of the other parents.

**Figure 3.**
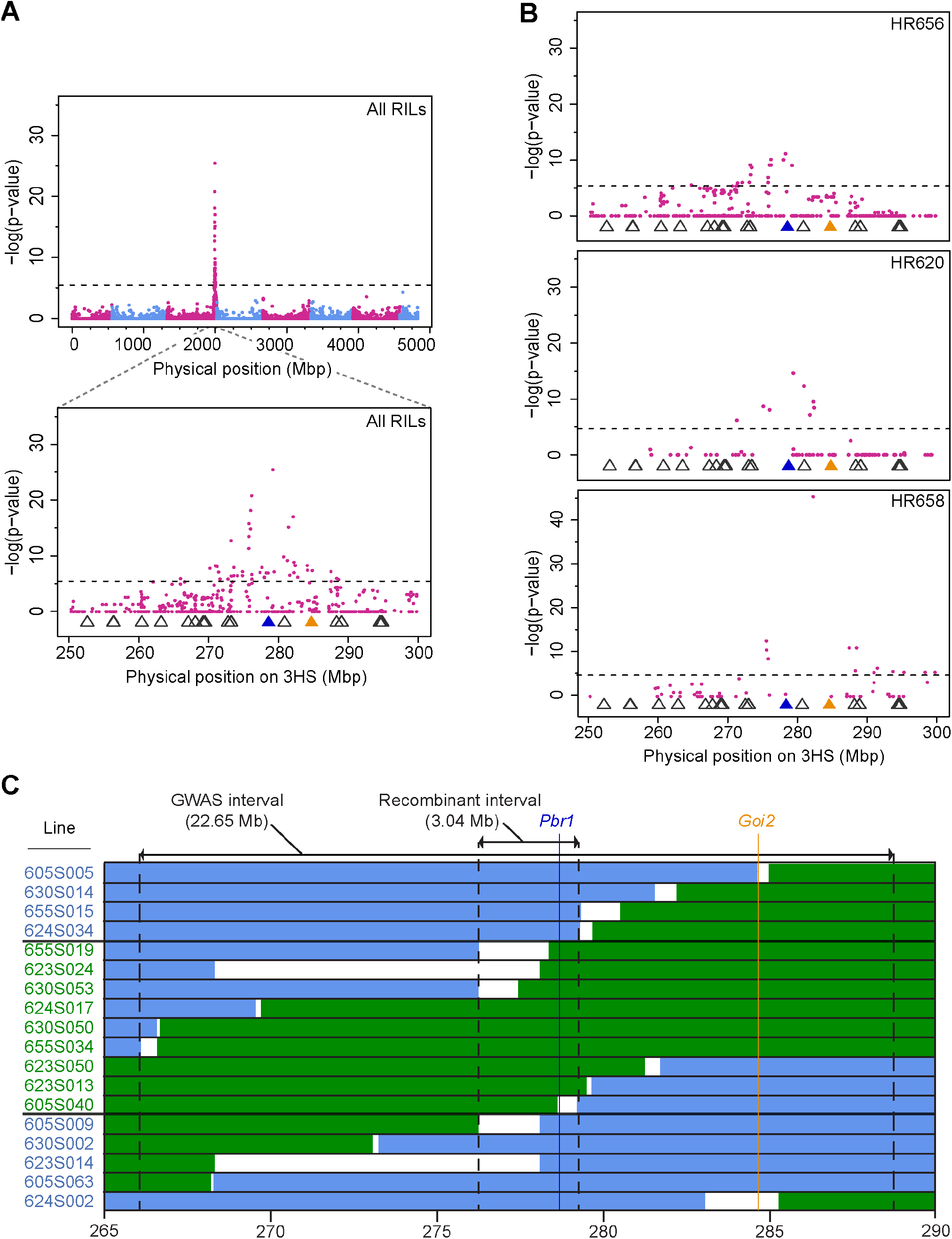
Genome wide association study identifies a single locus in the barley genome significantly associated with AvrPphB response. Manhattan plots of the association between SNPs and AvrPphB response of NAM barley lines for **A**) all 175 lines and **B**) the lines from each HR subpopulation individually. The X-axis shows SNPs in the region graphed, either the whole genome or the interval containing the significant locus in the short arm of Chromosome 3H (3HS). The Y-axis shows the negative logarithm of the p-value for the association. The locations of genes encoding NLRs predicted by NLR-Parser are indicated by open triangles; the blue triangle points to *Pbr1*, the orange to *Goi2*. The dotted horizontal line indicates a false discovery rate of 0.05 with Bonferroni correction. **C**) Graphical representation of 18 recombinant lines from four additional families used to fine map the AvrPphB response determinant. Green indicates regions containing SNPs matching the Rasmusson genotype. Blue indicates regions matching the other parental genotype. Uncolored regions represent the intervals in which it can be concluded the recombination took place, based on the nearest flanking SNPs. Lines labeled in green font display the Rasmusson HR phenotype (1), in blue the other parent phenotype (0).

Within the GWAS interval, there are 13 predicted NLR genes, as called by NLR-parser (Steuernagel et al., 2015) (Fig. 3A and 3B). In the reference genome, only four encode putative full length NLRs; the rest are fragments, mostly LRR domains and some partial NB-ARC domains. The most significant SNP in the analysis of all lines was S3H2_279293442 (3H:673604075; −log(p)=25.48). We selected the nearest predicted NLR to this SNP, HORVU3Hr1G107310 (3H: 672,928,614-672,932,121), as our top candidate for the determinant of the response to AvrPphB and tentatively named it *Pbr1* (*AvrPphB Response* 1). Despite its association with the most significant SNP, the protein product PBR1 is less closely related to RPS5 in both the encoded CC (Fig. 4A) and NB-ARC (Fig. 4B) domains than the protein products of the other NLR genes within the 22.65 Mb region. Of these, we selected the one encoding the NLR most closely related to RPS5, HORVU3Hr1G109680 (3H: 679064240-679072712), on the edge of the GWAS interval, as an additional gene of interest and refer to it hereafter as *Goi2*. PBR1 and RPS5 are 17% identical to each other at the CC domain and 29% at the NB-ARC domain, while GOI2 and RPS5 are 24% and 45% identical to each other at those domains, respectively. For comparison, PBR1 and GOI2 are 23% identical to each other at the CC domain and 20% at the NB-ARC domain.

**Figure 4.**
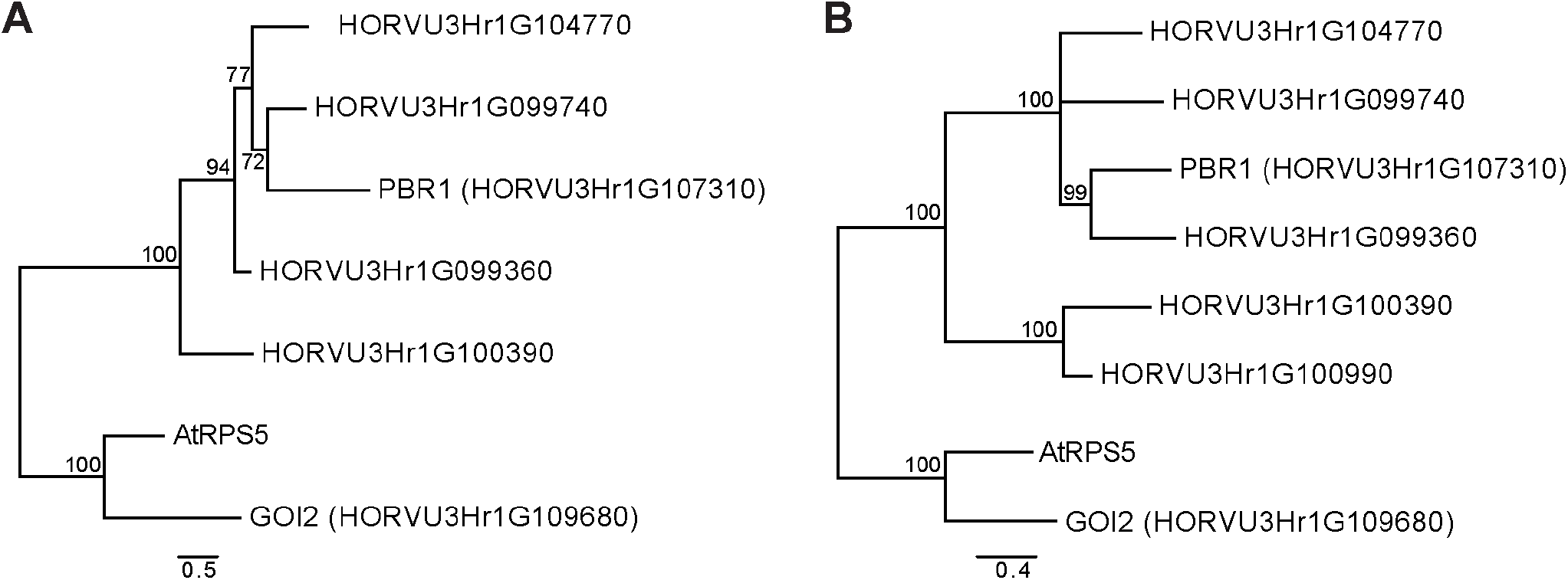
GOI2 is more closely related to RPS5 than PBR1 is, in both the CC and NB-ARC domains. Bayesian phylogenetic analysis of the **A**) coiled-Coil (CC) and **B**) nucleotide binding (NB-ARC) domains of Arabidopsis RPS5 (AtRPS5) and proteins encoded by predicted NLR genes in the GWAS interval on Chromosome 3HS. NB-ARC domains were extracted by NLR-Parser. CC domains were identified using the BLAST Conserved Domain search or by comparing to RPS5 for CC domains lacking the EDVID motif. Not all predicted NLR genes in the region encoded CC or NB-ARC domains. Scale bars indicate number of amino acid substitutions per site, and nodes are labeled with Bayesian posterior probabilities as a percentage. Albeit closest in the tree, GOI2 and RPS5 are only 24% identical at the CC domain and 45% at the NB-ARC domain. In comparison, PBR1 and RPS5 share 17% and 29% identity and PBR1 and GOI2 23% and 20% identity at those respective domains.

As a next step to identify the determinant of the AvrPphB response, we used SNP data for the entire NAM population to find additional recombinants within the 22.65 Mb GWAS interval. Based on haplotype data within the region, eighteen apparent recombinants were selected from four additional families with non-AvrPphB-responding parents and phenotyped (Supp. Table S2). Adding the genotype and phenotype data of these new lines to the GWAS increased the significance of many of the SNPs, but did not narrow the interval. However, using the estimated recombination breakpoints and the phenotypes of the individual RILs to fine map the determinant of the response resulted in a 3.04 Mb region within the GWAS peak that contains *Pbr1* and no other NLR gene (Fig. 3C), supporting *Pbr1* rather than *Goi2* as the candidate determinant.

### *Pbr1* is expressed in lines responding to AvrPphB and allelic variation correlates with phenotype

The reference genome used in the GWAS is from the barley line Morex, an AvrPphB-non-responding line (Fig. 1) (Mascher et al., 2017). Therefore, the reference genome is likely to have a nonfunctional copy of, or lack completely, the NLR hypothesized to detect activity of AvrPphB. In the Morex genome, *Pbr1* is annotated as containing just a truncated NB-ARC domain and LRR domain, missing an N-terminal domain (Marchler-Bauer and Bryant, 2004). In contrast, *Goi2* encodes a full length NLR (965 aa) with an RPS5-like CC domain (aa 27-66), NB-ARC domain (aa 156-439), and LRR (aa 537-864). To see if either gene sequence varies in the responding line Rasmusson, we sequenced *Pbr1* and *Goi2* from that line. The Rasmusson allele of *Goi2* is highly similar to the Morex allele, with only 3 nonsynonymous mutations between them (N860I, R808H, and V282L). Among the differences between the two *Pbr1* sequences, we found a single nucleotide insertion in Rasmusson that restores a larger open reading frame (Fig. 5A), resulting in a predicted full-length NLR (939 aa) with an intact CC_EDVID_ domain (aa 7-131), NB-ARC domain (aa 174-454), and an LRR domain containing 12 repeats (aa 474-886). For *Pbr1*, we will refer to the allele in Morex as *Pbr1.a* (GenBank: MH595617) and in Rasmusson as *Pbr1.b* (GenBank:MH595618).

**Figure 5.**
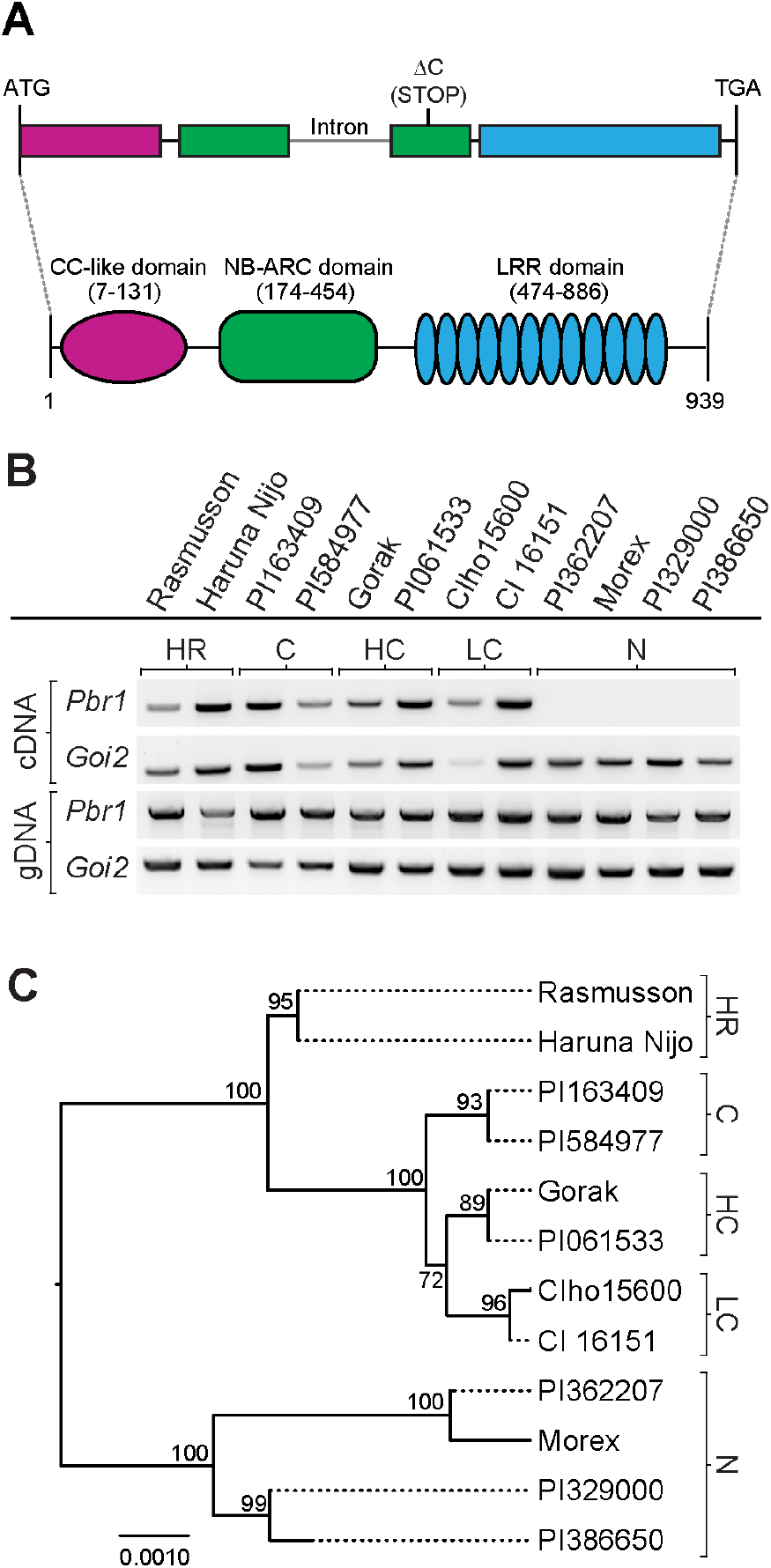
Sequence and expression polymorphism in *Pbr1* across barley lines correlate to AvrPphB response. **A**) Schematic illustration of *Pbr1.b*, the allele in the AvrPphB-responding line Rasmusson, showing the approximate location of a C nucleotide deletion that disrupts the open reading frame in *Pbr1.a*, the allele in the non-responding line Morex. Below, the *Pbr1.b* protein product is represented, with the amino acid positions of the CC domain, NB-ARC domain, and LRR domain indicated. **B**) PCR amplification from cDNA and genomic DNA (gDNA) of 12 representative lines that differ in their response to AvrPphB, showing expression and primer compatibility, respectively, for *Pbr1* and *Goi2*. cDNA was generated from RNA extracted from 10-day old plants, the same age used for phenotyping in Figure 1. See Figure S3 for data from additional lines. **C**) A neighbor joining tree showing the sequence relationships of *Pbr1* alleles from the barely lines represented in B) and the (at right) the responses of those lines to AvrPphB. The tree is based on aligned genomic DNA sequence from start codon to stop codon. Nodes are labeled with bootstrap values and the scale bar represents number of base substitutions per site. N, no response; LC, low chlorosis; C, chlorosis; HC, high chlorosis; HR, hypersensitive reaction.

Reverse transcriptase PCR (RT-PCR) was used to test the expression of *Pbr1* alleles in Morex and Rasmusson, as well as a variety of other barley lines ranging in AvrPphB-induced responses. *Pbr1* was expressed in lines that respond to AvrPphB either with HR or chlorosis (Rasmusson, Haruna Nijo, PI061533, Gorak, PI584977, PI163409, CIho15600, and CI 16151), but not in non-responding lines (PI329000, PI386650, PI362207, and Morex) (Fig. 5B). The primers used for RT-PCR were compatible with all genotypes tested, as shown by amplification from genomic DNA, and spanned an intron to differentiate cDNA from any genomic DNA contamination. We expanded our testing to 30 total lines: 12 responders, 12 non-responders, and 6 RILS from the 3 NAM subpopulations used for GWAS. *Pbr1* was expressed in all responding lines, but not expressed in 9 out of 12 non-responding lines (Fig. 5B and Supp. Fig. S3). For comparison, we assayed *Goi2* expression in these lines as well and found varying levels of expression that did not correspond to AvrPphB response (Fig. 5B).

Since point mutations within an NLR can lead to changes in observable HR *in planta* (Stirnweis et al., 2014), we were interested to see if the responses to AvrPphB that we observed across different barley lines corresponded with sequence polymorphism at *Pbr1*. We sequenced *Pbr1* alleles of 10 additional barley lines selected at random from among the different response phenotypes (1 HR, 2 HC, 2 C, 2 LC, and 3 nonresponding lines) and compared them to *Pbr1.a* and *Pbr1.b* from Morex and Rasmusson, respectively. The nucleotide sequences cluster by phenotype (Fig. 5C, Supp. File S1) when analyzed from start codon to stop codon using the Neighbor-Joining method. The non-responding lines, PI329000, PI386650, PI362207, and Morex have unique but similar alleles. The HR line Haruna Nijo, like Rasmusson, has the *Pbr1.b* allele. The amino acid sequences of *Pbr1* in the two lines each from the three chlorosis response groups (LC lines CI 16151 and CIho15600, C lines PI584977 and PI163409, and HC lines Gorak and PI061533) are identical within and different across the groups; all contain 3 common substitutions compared to the Rasmusson allele *Pbr1.b*, including an L538Q substitution in the LRR. Together these observations suggest that sequence polymorphism in *Pbr1* determines response to AvrPphB.

### The product of *Pbr1* allele *Pbr1.c* recognizes AvrPphB protease activity in *N. benthamiana*

To directly test whether PBR1 mediates recognition of AvrPphB, we developed a transient expression assay in *N. benthamiana. Pbr1.b* from cultivar Rasmusson was cloned into a dexamethasone-inducible vector along with a C-terminal fusion to super yellow fluorescent protein (PBR1.b.sYFP). Unfortunately, transient expression of PBR1.b:sYFP alone resulted in HR with complete tissue collapse within 24 hours of transgene induction (Fig. 6B), indicating that PBR1.b is auto-active when overexpressed in *N. benthamiana*.

**Figure 6.**
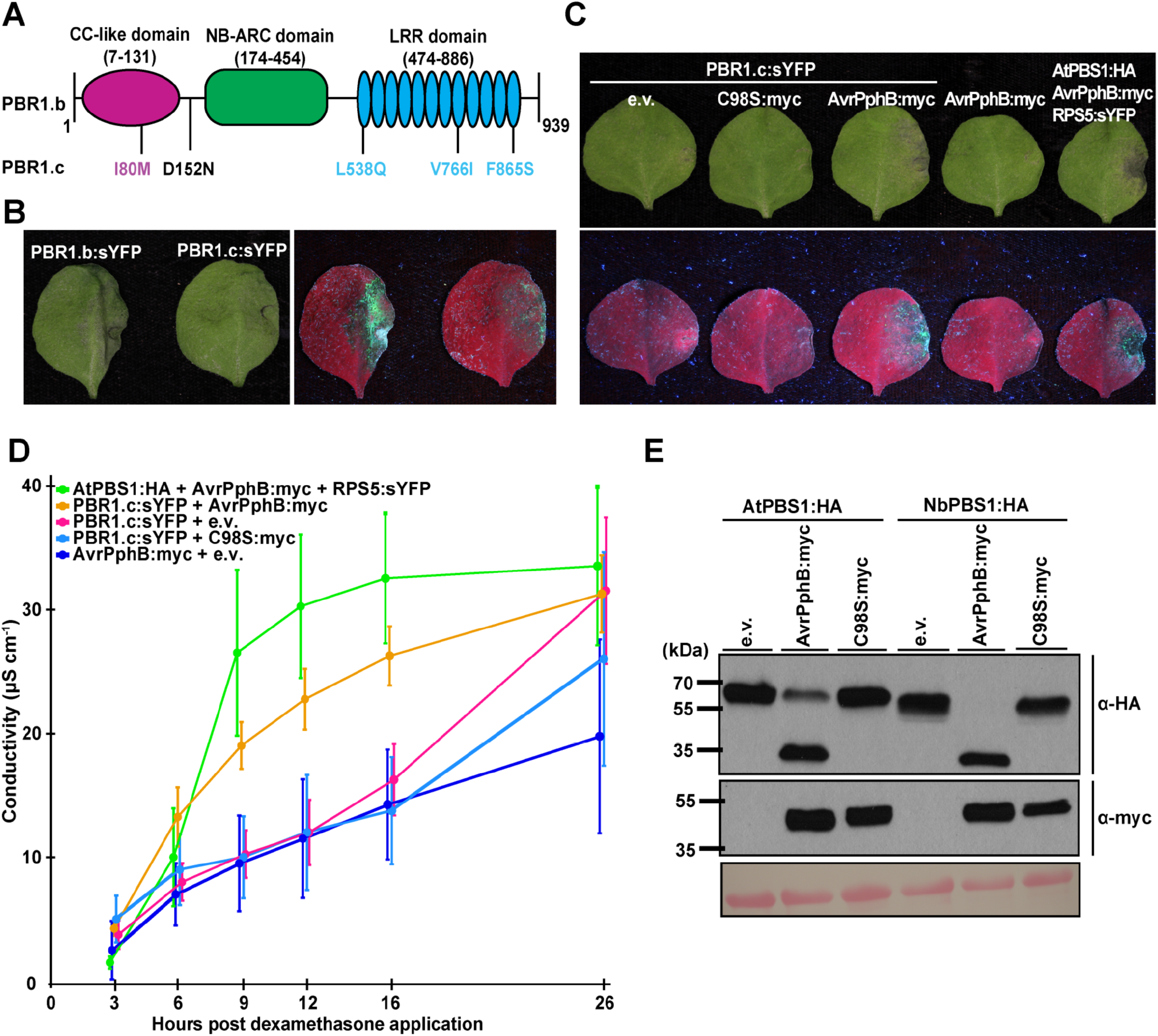
Transient co-expression of PBR1.c with AvrPphB induces cell death in *N. benthamiana*. **A**) Schematic representation of the PBR1.b protein product from Rasmusson, an HR line, showing the approximate locations of amino acid substitutions between the PBR1.b protein product and the PBR1.c protein product from CI 16151, a low-chlorosis line. **B**) Induction of cell death by PBR1.b:sYFP, but not PBR1.c:sYFP, independent of AvrPphB expression when transiently expressed in *N. benthamiana*. PBR1.b:sYFP or PBR1.c:sYFP were agroinfiltrated into 3-week old *N. benthamiana*. All transgenes were under the control of a dexamethasone-inducible promoter. A representative leaf was photographed 24 hours post-transgene induction under white light and UV light. Three independent experiments were performed with similar results. **C**) Activation of HR by transient co-expression of PBR1.c:sYFP and AvrPphB:myc in *N. benthamiana*. Agroinfiltrations were used to transiently express combinations of PBR1.c:sYFP, empty vector (e.v.), AvrPphB:myc, and a protease inactive derivative, AvrPphB(C98S):myc. HA-tagged Arabidopsis PBS1 coexpressed with RPS5:sYFP and AvrPphB:myc was used as a positive control. All transgenes were under the control of a dexamethasone-inducible promoter. A representative leaf was photographed 24 hours post-transgene induction under white light and UV light. Three independent experiments were performed with similar results. **D**) Electrolyte leakage as a measure of cell death resulting from co-expression of PBR1.c:sYFP with AvrPphB:myc relative to PBR1.c:sYFP with e.v. or AvrPphB(C98S):myc in *N. benthamiana* leaf discs. Leaf discs were transiently expressing the indicated combinations of constructs. Conductivity is shown as mean ± S.D. (n = 4). Three independent experiments were performed with similar results. **E**) Cleavage of *N. benthamiana* PBS1 (NbPBS1) by AvrPphB. HA-tagged *N. benthamiana* PBS1 or AtPBS1 was transiently co-expressed with or without myc-tagged AvrPphB or AvrPphB(C98S) in *N. benthamiana*. Total protein was extracted six hours post-transgene induction and immunoblotted with the indicated antibodies. Two independent experiments were performed with similar results.

To circumvent the problem posed by auto-activity of the PBR1.b protein, we tested a *Pbr1* allele from the LC line CI 16151 (Fig. 1). We designated this allele *Pbr1.c* (GenBank: MH595619). PBR1.b and PBR1.c differ by five amino acid substitutions, of which three are located within the leucine-rich repeat domain (Fig 6A). Transient expression of a PBR1.c:sYFP fusion protein in the absence of AvrPphB consistently produced a weaker HR than PBR1.b (Fig. 6B). This result allowed us to test whether the HR was enhanced in the presence of active AvrPphB.

We transiently co-expressed PBR1.c:sYFP and AvrPphB:myc in *N. benthamiana* and assessed cell death. As a control, we co-expressed AtPBS1:HA and RPS5:sYFP with AvrPphB:myc, a combination that activates cell death in *N. benthamiana* (Ade et al., 2007; DeYoung et al., 2012; Qi et al., 2014). Transient co-expression of PBR1.c:sYFP with AvrPphB:myc resulted in observable tissue collapse 24 hours post-transgene induction, whereas co-expression of PBR1.c:sYFP with either empty vector (e.v.) or AvrPphB(C98S):myc resulted in a much weaker cell death response (Fig. 6C). Further, transient expression of AvrPphB:myc in the absence of PBR1.c:sYFP did not trigger HR, indicating that the cell death response requires PBR1.c (Fig. 6C). We performed an electrolyte leakage analysis to better quantify PBR1.c-mediated cell death. Transient co-expression of PBR1.c:sYFP with AvrPphB-myc induced greater ion leakage than PBR1.c:sYFP co-expressed with either empty vector or AvrPphB(C98S):myc between 9 and 16 hours after transgene induction, confirming that PBR1.c:sYFP recognizes and mediates a response to AvrPphB protease activity (Fig. 6D). By 26 hours post transgene induction, PBR1.c:sYFP expressed with AvrPphB(C98S) or empty vector induced ion leakage similar to that observed with co-expression of PBR1.c:sYFP and wild-type AvrPphB, indicating that PBR1.c:sYFP is weakly auto-active, consistent with the HR assays (Fig. 6D).

The observation that AvrPphB, but not AvrPphB(C98S), activates PBR1.c-mediated cell death in *N. benthamiana* even in the absence of a barley PBS1 protein suggested that AvrPphB might be cleaving an *N. benthamiana* ortholog of PBS1 and that PBR1.c is recognizing that cleavage. Using a reciprocal BLAST and the amino acid sequence of Arabidopsis PBS1, we identified an ortholog of PBS1 in the *N. benthamiana* genome, Niben101Scf02996g03008.1 (Bombarely et al., 2012), and designated it *NbPBS1*. Importantly, NbPBS1 contains the AvrPphB cleavage site sequence and is thus predicted to be cleaved by AvrPphB. To test our hypothesis that in the transient assay PBR1.c is guarding an endogenous PBS1 ortholog, we coexpressed NbPBS1:HA with either AvrPphB:myc or AvrPphB(C98S):myc. Consistent with our hypothesis, co-expression with AvrPphB:myc, but not the protease inactive mutant, resulted in cleavage of NbPBS1:HA within 6 hours post-transgene expression, showing that NbPBS1:HA is a substrate for AvrPphB (Fig. 6E) and that its cleavage could be the trigger for PBR1.c.

### PBS1 proteins immunoprecipitate with barley PBR1.c when transiently co-expressed in *N. benthamiana*

To further test the hypothesis that PBR1.c is activated by sensing cleavage of PBS1 proteins, we performed co-immunoprecipitation (co-IP) analyses of PBR1.c with HvPBS1-1:HA, HvPBS1-2:HA, AtPBS1:HA, or NbPBS1:HA. As a positive control, we co-expressed AtPBS1:HA with RPS5:sYFP, which forms a preactivation complex in the absence of AvrPphB (Ade et al., 2007). As a negative control, we transiently coexpressed the plasma membrane-localized fusion protein sYFP:LTI6b with each of the PBS1 proteins (Cutler et al., 2000). Consistent with our hypothesis, HvPBS1-1:HA, HvPBS1-2:HA, and AtPBS1:HA immunoprecipitated with PBR1.c:sYFP and not with sYFP:LTI6b, demonstrating that PBR1.c forms a complex with PBS1 proteins from barley and Arabidopsis in the absence of AvrPphB (Fig. 7). NbPBS1:HA also immunoprecipitated with PBR1.c:sYFP (and not with sYFP:LTI6b) supporting the notion that AvrPphB-mediated cleavage of NbPBS1 activates PBR1.c-dependent HR in *N. benthamiana*. Though all of the PBS1 proteins immunoprecipitated with PBR1.c:sYFP, PBR1.c:sYFP preferentially interacted with HvPBS1-2:HA and AtPBS1:HA (Fig. 7). Collectively, these data suggest that PBR1 forms a pre-activation complex with one or more barley PBS1 orthologs, providing further evidence that PBR1 is the guard that recognizes AvrPphB activity. Importantly, CSS-PALM 4.0 (http://csspalm.biocuckoo.org/) predicts that PBR1.b and PBR1.c are palmitoylated at Cys314, suggesting co-localization with AvrPphB and barley PBS1 orthologs at the plasma membrane (Ren et al., 2008; Dowen et al., 2009; Sun et al., 2017).

**Figure 7.**
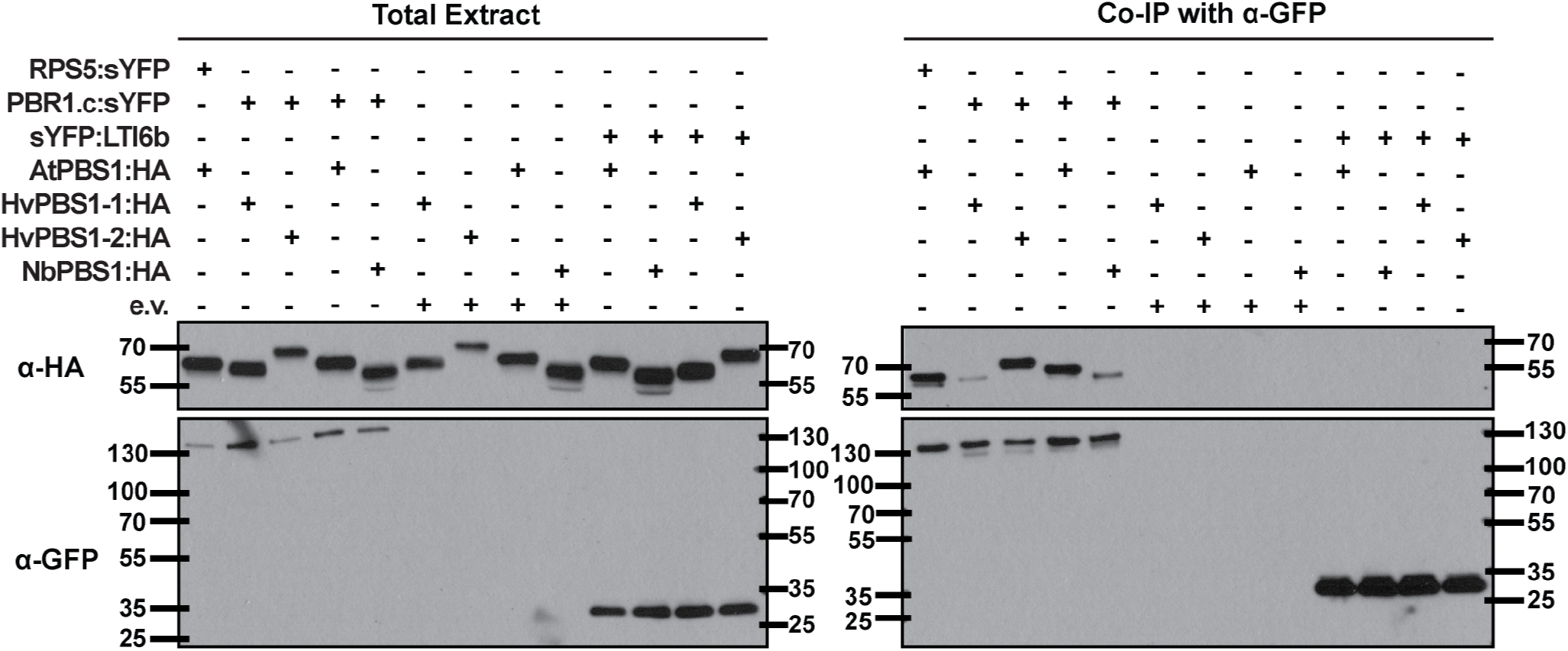
PBS1 proteins immunoprecipitate with PBR1.c when transiently co-expressed in *N. benthamiana*. The indicated construct combinations were transiently co-expressed in leaves of 3-week old *N. benthamiana* plants using agroinfiltration. All transgenes were under the control of a dexamethasone-inducible promoter. Total protein was extracted six hours post-transgene induction. HA-tagged Arabidopsis PBS1 coexpressed with RPS5:sYFP was used as a positive control. The sYFP:LTI6b fusion protein, which is targeted to the plasma membrane (Cutler et al., 2000), was co-expressed with the HA-tagged PBS1 proteins as a negative control. Results are representative of two independent experiments.

### Wheat (*Triticum aestivum* subsp. *aestivum*) also recognizes AvrPphB protease activity

Sun et al. (2017) recently identified an ortholog of Arabidopsis PBS1 in wheat, TaPBS1, that localizes to the plasma membrane when transiently expressed in *N. benthamiana* and is cleaved by AvrPphB. However, it remained unclear whether wheat recognizes AvrPphB protease activity and would thus likely contain a functional analog of *RPS5*, such as *Pbr1*. We screened 34 wheat varieties obtained from the U.S. Department of Agriculture Wheat Germplasm Collection for their response to D36E expressing AvrPphB (Fig. 8A; Supp. Table S3). Twenty-nine responded with chlorosis, while five showed no visible response by three days post-inoculation (Fig. 8A; Supp. Table S3). No line responded to the protease inactive mutant AvrPphB(C98S).

**Figure 8.**
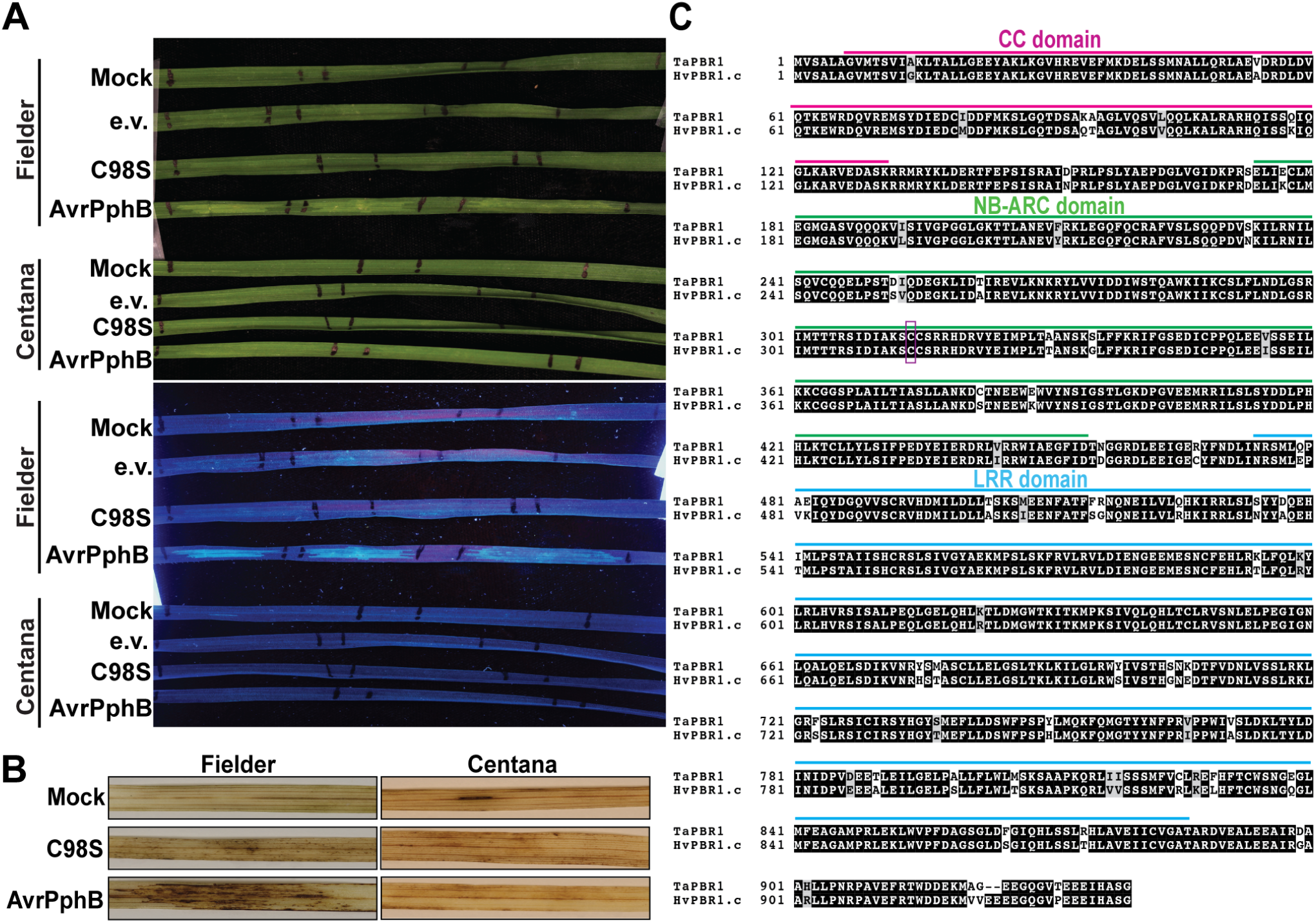
Recognition of AvrPphB protease activity is conserved in wheat. **A**) Responses of wheat cultivars Fielder and Centana infiltrated with (top to bottom) 10 mM MgCl_2_ (mock), *P. syringae* DC3000(D36E) expressing empty vector (e.v.), AvrPphB(C98S), or AvrPphB three days post-infiltration (dpi), photographed under white and UV light. Bacteria (OD_600_= 0.5) were infiltrated into the adaxial surface of the second leaf of two-week old seedlings. Three independent experiments were performed with similar results. Responses of all lines tested are recorded in Table S3. **B**) Hydrogen peroxide accumulation. Cultivars and treatments assayed were as in A). Three dpi, leaf segments were excised from the infiltrated regions, stained with DAB solution, cleared with 70% ethanol, and photographed under white light. This experiment was repeated twice with similar results. **C**) Full-length amino acid sequence alignment between barley PBR1.c and the most closely related homolog in wheat, TRIAE_CS42_3B_TGACv1_226949_AA0820360 (TaPBR1). Conserved residues and conservative substitutions are highlighted with black and grey backgrounds, respectively. The predicted coiled-coil (CC), nucleotide binding (NB-ARC), and leucine-rich repeat (LRR) domains of TaPBR1 are indicated by pink, green, and cyan bars, respectively. The predicted palmitoylation site is indicated by a purple box.

To further characterize the chlorotic response in wheat, we used 3,3’-diaminobenzidine (DAB) staining to examine hydrogen peroxide accumulation following leaf infiltration with D36E expressing either AvrPphB or AvrPphB(C98S). Consistent with the chlorotic phenotype, wheat cv. Fielder accumulated detectable hydrogen peroxide within the infiltrated area when inoculated with D36E expressing AvrPphB, whereas the mock and AvrPphB(C98S) treatments resulted in minimal hydrogen peroxide accumulation (Fig. 8B). In contrast, there was no significant hydrogen peroxide accumulation in wheat cv. Centana inoculated with either strain (or mock), consistent with the lack of chlorotic response of this line to AvrPphB. The correlation of chlorosis and hydrogen peroxide accumulation specifically in response to active AvrPphB is consistent with recognition in wheat associated with defense.

To examine whether this recognition in wheat might be mediated by a *Pbr1* ortholog, we searched the *T. aestivum* subsp. *aestivum* genome using the Ensembl genome browser (release TGACv1) (Clavijo et al., 2017; Kersey et al., 2018). We found TRIAE_CS42_3B_TGACv1_226949_AA0820360 to be an ortholog, and designated it *TaPbr1. TaPbr1* is located on wheat chromosome 3B in a position syntenic with barley *Pbr1* and encodes an NLR consisting of a predicted Rx-like coiled-coil domain (aa 7-131), a nucleotide-binding domain (aa 174-454), and a leucine-rich repeat domain (aa 474-895) (Fig. 8C). Full-length amino acid sequence alignment of barley PBR1.c and TaPBR1 shows 93% amino acid identity (Fig. 8C). Further, TaPBR1, like barley PBR1, is predicted to be palmitoylated at Cys314, suggesting co-localization with AvrPphB and wheat PBS1. It thus seems likely that TaPBR1 functions as the cognate NLR protein that mediates recognition of AvrPphB protease activity in wheat.

## Discussion

Recognition of the *P. syringae* AvrPphB protease by the Arabidopsis RPS5 NLR protein is a well characterized example of indirect effector recognition (Kim et al., 2016). Though AvrPphB is recognized by other plant species such as soybean and common bean, the disease resistance genes responsible for recognition outside of Arabidopsis have not been cloned, and the underlying molecular mechanisms are unknown (Jenner et al., 1991; Russell et al., 2015). The evidence herein supports the conclusion that barley and Arabidopsis have convergently evolved NLRs able to detect effectors that structurally modify PBS1-like kinases: barley cultivars respond to AvrPphB but not to a protease inactive mutant of AvrPphB, barley contains an NLR gene evolutionarily distinct from RPS5 that mediates a strong HR when co-expressed with *avrPphB* in *N. benthamiana*, and AvrPphB associates with and cleaves PBS1 orthologs from monocots and dicots.

While AvrPphB is not known to be present in any pathogens of barley, it is a member of a family of proteases present in many phytopathogenic bacteria (Shao et al., 2002; Dowen et al., 2009). More generally, proteases that target host proteins are found in many, diverse types of pathogens, and we expect conserved kinases that are involved in PTI to be common effector targets (Xia, 2004). Though the functional roles of HvPBS1-1 and HvPBS1-2 as well as other barley PBS1-like proteins are unknown, given their conservation in many flowering plant families, we can predict that they have a role in PTI signaling as observed in Arabidopsis (Zhang et al., 2010). Our evidence supports the hypothesis that barley deploys an effector protease recognition mechanism similar to that of recognition of AvrPphB by Arabidopsis RPS5, wherein barley PBR1 guards RLCKs such as HvPBS1-1 and HvPBS1-2 such that it is activated upon their cleavage. Within Arabidopsis populations, RPS5 is maintained as a balanced presence/absence polymorphism despite inconsistent interaction with *Pseudomonas* strains expressing AvrPphB homologs, suggesting other effectors are also imposing selection pressure (Tian et al., 2002; Karasov et al., 2014). How many and which effectors from barley pathogens target RLCKs is unknown.

The convergent evolution of the shared ability of PBR1 and RPS5 to recognize AvrPphB aligns with the prediction that RLCKs that function in plant immunity are common targets of pathogen effectors and that selection to guard these proteins is ancient and widespread. A similar example of convergent evolution of NLR specificity has been described for the RPM1 and Rpg1b/Rpg1r proteins of Arabidopsis and soybean, all three of which detect effector induced modifications of RIN4 proteins (Ashfield et al., 2004; Selote and Kachroo, 2010; Ashfield et al., 2014). Like PBL proteins, RIN4 is targeted by multiple effectors, consistent with these proteins serving critical functions in plant immunity (Afzal et al., 2013). It is especially interesting that PBR1 and RPS5 independently evolved to detect PBL cleavage instead of directly interacting with AvrPphB or integrating a PBL decoy. Direct interaction limits the number of effectors a single NLR can detect, while guarding a commonly targeted host protein expands the response spectrum, thus allowing the NLR to detect multiple pathogen effectors. The guarding strategy might impose purifying selection on RLCKs themselves or selection to integrate an RLCK decoy into an NLR: either would reduce the risk of any guard-guardee genetic mismatch that might lead to hybrid necrosis. However, there is no obvious reason why PBR1 and RPS5 would both have each evolved to guard PBS1, rather than distinct avrPphB substrates, and our current data do not rule out the possibility that in barley PBR1 is activated by cleavage of one or more different RLCKs.

When assessing the functional role of PBR1.c in AvrPphB recognition, we showed that its co-expression with AvrPphB elicited cell death in *N. benthamiana* even in the absence of barley PBS1 expression. Given that PBR1.c does not contain the AvrPphB cleavage site sequence, it is likely PBR1.c is sensing AvrPphB-mediated cleavage of an endogenous *N. benthamiana* PBL protein, and we showed that the PBS1 ortholog NbPBS1 indeed associates with PBR1 in the absence of AvrPphB and is cleaved by AvrPphB. This was true also of AtPBS1 and the barley PBS orthologs HvPBS1-1 and HvPBS1-2. Taken together, these data strongly suggest that in barley, PBR1.c detects AvrPphB protease activity by sensing cleavage of a PBS1 protein, analogous to AvrPphB-detection by RPS5.

*Pbr1* is expressed in the 12 tested barley lines that respond to AvrPphB, and in only 3 of 12 lines that do not respond. The sequence polymorphisms found in *Pbr1* alleles across the 12 responding barley lines correlate with the presence and severity of the AvrPphB response (i.e. chlorosis versus strong HR). These data suggest that mutations within the *Pbr1* coding sequence impact the macroscopic phenotype observed when AvrPphB is present. Mutagenesis screens of specific NLRs have been shown to modify the severity of phenotype and specificity of interaction (Farnham and Baulcombe, 2006; Harris et al., 2013; Segretin et al., 2014). Natural examples of the effect of single or few mutations impacting NLR function include the Pi-ta NLR in *Oryzae* spp., in which a single amino acid is highly correlated to resistance, and the barley *Mla* locus, which encodes alleles with over 90% amino acid sequence identity that recognize different effector proteins (Huang et al., 2008; Lu et al., 2016). In wheat, alleles of the *Pm3* gene have very little sequence diversity, but just two amino acid mutations expand the effector recognition capacity of *Pm3f* and increase its activity (Brunner et al., 2010; Stirnweis et al., 2014). We have not yet functionally characterized the polymorphisms in PBR1 to determine which, if any, modify the response to AvrPphB response or if any impact specificity. However, the difference in auto-activity between PBR1.b and PBR1.c when expressed in *N. benthamiana* is further evidence that the allelic sequence polymorphism contributes to phenotype.

The evidence that PBR1 is activated by cleavage of a PBS1 or PBS1-like protein suggests that PBS1-based decoys can be used to expand protease effector recognition in barley. Barley powdery mildew (*Blumeria graminis* f. sp. *hordei; Bgh*) and *Wheat streak mosaic virus* (WSMV) are two barley pathogens known to deploy proteases as part of the infection process (Pliego et al., 2013; Singh et al., 2018). BEC1019 is a putative metalloprotease secreted by *Bgh* and is conserved among ascomycete fungi (Pliego et al., 2013; Whigham et al., 2015). Notably, silencing of BEC1019 by both *Barley stripe mosaic virus*- and single cell RNAi-based methods reduces *Bgh* virulence, suggesting BEC1019 is required for *Bgh* pathogenicity (Pliego et al., 2013; Whigham et al., 2015). Similar to other *Potyviruses*, WSMV expresses a protease, designated the nuclear inclusion antigen (NIa), that is essential for viral replication and for proper temporal expression of potyviral genes *in planta* (Singh et al., 2018). Importantly, the cleavage site sequence recognized by the NIa protease has been identified (Choi et al., 2001; Tatineni et al., 2011). Insertion of the BEC1019 or NIa protease cleavage site sequence into the barley PBS1 proteins should enable recognition of these proteases by PBR1. This approach could also be extended into wheat given that PBR1 and PBS1 are conserved.

## Materials and Methods

### Plant Material and Growth Conditions

Barley seeds were planted in Cornell mix soil (1.2 cubic yards of mix contains 10.6 cubic feet of compressed peat moss, 20 lb of dolomitric limestone, 6 lb of 11-5-11 fertilizer, 12 cubic ft of vermiculite) in plastic pots. Barley plants were grown in a growth room on a 16 hr light/8 hr dark cycle with cool white fluorescent lights (85 to 112 μmol/m^2^/s at soil level) at 22°C. Plants were watered as needed to keep soil damp.

*N. benthamiana* seeds were sown in plastic pots containing Pro-Mix B Biofungicide potting mix supplemented with Osmocote slow-release fertilizer (14-14-14) and grown under a 12 hr photoperiod at 22°C in growth rooms with average light intensities at plant height of 150 μEinsteins/m^2^/s.

Seed for wheat (*Triticum aestivum* subsp. *aestivum*) cultivars were ordered from the U.S. Department of Agriculture Wheat Germplasm Collection via the National Plant Germplasm System Web portal (https://www.ars-grin.gov/npgs/) or provided by S. Hulbert (Washington State University). Wheat plants were grown in clay pots containing Pro-Mix B Biofungicide potting mix supplemented with Osmocote slow-release fertilizer (14-14-14) and grown under a 12 hr photoperiod at 22°C in growth rooms with average light intensities at plant height of 150 μEinsteins/m^2^/s.

### *P. syringae* DC3000(D36E) *in planta* assays

Previously generated plasmids pVSP61-AvrPphB and pVSP61-AvrPphB(C98S) (a catalytically inactive mutant) (Simonich and Innes, 1995; Shao et al., 2003) were each transformed into D36E, a strain of *Pseudomonas syringae* pv. *tomato* DC3000 with 36 effectors removed (Wei et al., 2015). Bacteria were grown on King’s media B (KB), supplemented with 50 μg of kanamycin per milliliter, for two days at 28°C, then suspended in 10 mM MgCl_2_ to an OD_600_ of 0.5. Suspensions were infiltrated into the underside of the primary leaf of 10-day old barley seedlings by needleless syringe. Each leaf was infiltrated with bacteria expressing AvrPphB and bacteria expressing AvrPphB(C98S), and the infiltrated areas were marked with permanent marker. Infiltrated leaves were checked for cell collapse two days post infiltrations, then photographed and phenotyped for chlorosis and necrosis five days post infiltrations.

For wheat inoculations, bacteria were grown and prepared in the same way, but the adaxial side of the second leaf of 14-day old wheat seedlings was infiltrated at three spots with one of the strains of bacteria per leaf. Responses were photographed three days after infiltration using a high intensity long-wave (365 nm) ultraviolet lamp (Black-Ray B-100AP, UVP, Upland, CA).

### Phylogenetic Analyses

Homology searches were performed using BLASTp to gather barley amino acid sequences homologous to Arabidopsis PBS1 and PBS1-like proteins. First, AtPBL (1 to 27), BIK1, and other PBS1-homologous sequences were gathered by searching the Arabidopsis genome (TAIR10, GCA_000001735.1) with the AtPBS1 (OAO91748.1) amino acid sequence and by name search. Potential barley PBLs were collected by searching the barley protein database (assembly Hv_IBSC_PGSB_v2) with each Arabidopsis homologue and taking the top five hits derived from distinct genes.

For NLR phylogenetic analysis, the NB-ARC domain was extracted by NLR-parser (Steuernagel et al., 2015). For genes where no NB-ARC domain was automatically found, the upstream nucleotide sequence in the genome was inspected using BLASTx to look for fragments encoding an NB-ARC domain or CC domain. CC domains were identified by analyzing each predicted NLR with the BLAST Conserved Domain Search or by comparison to the CC domain in RPS5 for domains lacking the EDVID motif (Marchler-Bauer and Bryant, 2004).

Nucleotide or amino acid sequences were aligned with Clustal Omega (Sievers et al., 2011). Bayesian phylogenetic trees were generated for the collected sequences using the program MrBayes under a mixed amino acid model (Ronquist et al., 2012). Parameters for the Markov chain Monte Carlo method were; nruns = 2, nchains = 2, diagnfreq = 1000, diagnstat = maxstddev. The number of generations (ngen) was initially set at 200,000 and increased by 100,000 until the max standard deviation of split frequencies was below 0.01, or until it was below 0.05 after 1,000,000 generations. Phylogenetic trees were visualized in FigTree v1.4.3.

For the analysis of *Pbr1* alleles, nucleotide sequences were selected from each sequenced allele that spanned from the start codon to the stop codon of the Rasmusson allele, including the intron. Sequences were aligned with Clustal Omega and then used to construct Neighbor-Joining trees in MEGA7 (Kumar et al., 2016). A bootstrap test of 1000 replicates was applied.

### Genome Wide Association Study

The University of Minnesota Spring Barley Nested Association Mapping (NAM) population comprises 6,161 RILs generated from the variety Rasmusson crossed to 88 diverse parents that represent 99.7% of captured SNP diversity. In total, ~24,000 SNPs were generated through use of genotyping by sequencing and the barley iSelect 9K SNP chip. The 89 parental lines were assayed for AvrPphB response as part of the initial survey of barley lines. Because the common parent, Rasmusson, displayed a strong hypersensitive response, NAM families derived from Rasmusson and a parent showing no response were chosen for GWAS.

Plants were assayed as described above using infiltrations of two *Pseudomonas* strains expressing either AvrPphB or AvrPphB(C98S). Phenotypes for at least six plants of each recombinant inbred line (RIL) were recorded as 0 (no response/low chlorosis) or 1 (hypersensitive reaction) depending on the parental phenotype they exhibited. Lines that showed phenotypic segregation between individuals were not included in the analysis.

Genome wide association analysis was performed with the gwas2 function from the R/NAM (Nested Association Mapping) package, which uses an empirical Bayesian framework to determine likelihood ratios for each marker (Xavier et al., 2015). Lines from each family were identified within a family vector to account for population stratification. Markers with a minor allele frequency below 0.05 or missing data of more than 20% were removed using the snpQC function prior to analysis. A threshold of 0.05 for the false discovery rate was used to identify significant associations. NLR-encoding gene prediction was generated using NLR-parser (Steuernagel et al., 2015) and the high confidence Morex barley genome protein predictions (Mascher et al., 2017).

For genetic fine mapping, eighteen additional RILs with recombination events in the GWAS interval were selected from other families that also had an AvrPphB-non-responding parent. To determine which RILs to select, we subset the master SNP file by family and removed SNPs that were not variable between Rasmusson and the other parent. For visualization, SNPs that did not match neighboring markers across RILs were assumed to be miscalls and were also removed; while these could indicate double recombination events, the probability for a double recombination occurring within the 22.65 Mb interval is 0.001, and would be even less between two or three SNPs.

### Construction of Transgene Expression Plasmids

The AvrPphB:myc, AvrPphB(C98S):myc, RPS5:sYFP, and AtPBS1:HA constructs have been described previously (Shao et al., 2003; Ade et al., 2007; DeYoung et al., 2012). HORVU2Hr1G070690 (*HvPbs1-1*) and HORVU3Hr1G035810 (*HvPbs1-2*) were PCR amplified from barley accession CI 16151 (Manchuria background) and Rasmusson cDNA, respectively. The resulting fragments were gel-purified, using the QIAquick gel extraction kit (Qiagen), and cloned into the Gateway entry vector pCR8/GW/TOPO (Invitrogen) to generate pCR8/GW/TOPO:HORVU2Hr1G070690 and pCR8/GW/TOPO:HORVU3Hr1G035810, which we then designated pCR8/GW/TOPO:HvPbs1-1 and pCR8/GW/TOPO:HvPbs1-2, respectively.

The following genes were PCR amplified with attB-containing primers from the corresponding templates: *HvPbs1-1* from pCR8/GW/TOPO:*HvPbs1-1*, *HvPbs1-2* from pCR8/GW/TOPO:HvPbs1-2, *Pbr1.b* (HORVU3Hr1G107310) and *Goi2* (HORVU3Hr1G109680) from Rasmusson cDNA, *Pbr1.c* from CI 16151 gDNA, *LTI6b* from *Arabidopsis thaliana* gDNA (Col-0), and *NbPbs1* (Niben101Scf02996g03008.1) from *Nicotiana benthamiana* cDNA. The resulting PCR products were gel-purified, using the QIAquick gel extraction kit (Qiagen) or the Monarch DNA gel extraction kit (NEB), and recombined into the Gateway donor vectors pBSDONR(P1-P4) or pBSDONR(P4r-P2) using the BP Clonase II kit (Invitrogen) (Qi et al., 2012). The resulting constructs were sequence-verified to check for proper sequence and reading frame.

To generate protein fusions with the desired C-terminal epitope tags, pBSDONR(P1-P4):HvPbs1-1, pBSDONR(P1-P4):HvPbs1-2, and pBSDONR(P1-P4):NbPbs1 were mixed with the pBSDONR(P4r-P2):3xHA construct and the Gateway-compatible expression vector pBAV154 in a 2:2:1 molar ratio. A derivative of the destination vector pTA7001, pBAV154, carries the dexamethasone inducible promoter (Aoyama and Chua, 1997; Vinatzer et al., 2006). The pBSDONR(P1-P4):Pbr1.b and pBSDONR(P1-P4):Pbr1.c constructs were mixed with the pBSDONR(P4r-P2):sYFP construct and pBAV154 in a 2:2:1 molar ratio. The pBSDONR(P4r-P2):sYFP and pBSDONR(P4r-P2):3xHA constructs have been described previously (Qi et al., 2012). To generate the sYFP:LTI6b fusion protein, the pBSDONR(P4r-P2):LTI6b construct was mixed with the pBSDONR(P1-P4):sYFP construct and pBAV154 in a 2:2:1 molar ratio. Plasmids were recombined by the addition of LR Clonase II (Invitrogen) and incubated overnight at 25°C following the manufactures instructions. Constructs were sequence verified and subsequently used for transient expression assays in *N. benthamiana*.

### Transient Expression Assays in *N. benthamiana*

For transient expression assays in *N. benthamiana*, we followed the protocol described by DeYoung et al. (2012) and Kim et al. (2016). Briefly, the dexamethasone-inducible constructs were transformed into *Agrobacterium tumefaciens* GV3101 (pMP90) strains and were streaked onto Luria-Bertani (LB) plates containing 30 μg of gentamicin sulfate per milliliter and 50 μg of kanamycin per milliliter. Cultures were prepared in liquid LB media (5 ml) supplemented with 30 μg of gentamicin per milliliter and 50 μg of kanamycin per milliliter and shaken overnight at 30°C and 250 rpm on a New Brunswick orbital shaker. After overnight culture, the bacterial cells were pelleted by centrifuging at 3000 x g for 3 minutes and resuspended in 10 mM MgCl_2_ supplemented with 100 μM acetosyringone (Sigma-Aldrich). The bacterial suspensions were adjusted to an OD_600_ of 0.9 for HR and electrolyte leakage assays and an OD_600_ of 0.3 for immunoprecipitation and immunoblotting assays, and incubated for 3 hours at room temperature. For co-expression of multiple constructs, suspensions were mixed in equal ratios. Bacterial suspension mixtures were infiltrated by needleless syringe into expanding leaves of 3-week-old *N. benthamiana*. Leaves were sprayed with 50 μM dexamethasone 45 hours after injection to induce transgene expression. Samples were harvested 6 hours after dexamethasone application for protein extraction, flash-frozen in liquid nitrogen, and stored at −80°C. HR was evaluated and leaves photographed 24 hours after dexamethasone application using a high intensity longwave (365 nm) ultraviolet lamp (Black-Ray B-100AP, UVP, Upland, CA).

### Immunoblot Analysis

Frozen *N. benthamiana* leaf tissue (0.5 g) was ground in two volumes of protein extraction buffer (150 mM NaCl, 50 mM Tris [pH 7.5], 0.1% Nonidet P-40 [Sigma-Aldrich], 1% plant protease inhibitor cocktail [Sigma-Aldrich], and 1% 2,2’-dipyridyl disulfide [Chem-Impex]) using a ceramic mortar and pestle and centrifuged at 10,000 x g for 10 minutes at 4°C to pellet debris. Eighty microliters of total protein lysate were combined with 20 μl of 5X SDS loading buffer, and the mixture was boiled at 95°C for 10 minutes. All samples were loaded on a 4-20% gradient Precise™ Protein Gels (Thermo Fisher Scientific, Waltham, MA) and separated at 185 V for 1 hour in 1X Tris/Glycine/SDS running buffer. Total proteins were transferred to a nitrocellulose membrane (GE Water and Process Technologies, Trevose, PA). Ponceau staining was used to confirm equal loading of protein samples and successful transfer. Membranes were washed with 1X Tris-buffered saline (TBS; 50 mM Tris-HCl, 150 mM NaCl, pH 7.5) solution containing 0.1% Tween 20 (TBST) and blocked with 5% Difco™ Skim Milk (BD, Franklin Lakes, NJ) overnight at 4°C. Proteins were detected with 1:5,000 diluted peroxidase-conjugated anti-HA antibody (rat monoclonal, Roche, catalog number 12013819001) and a 1:5,000 diluted peroxidase-conjugated anti-c-Myc antibody (mouse monoclonal, Thermo Fisher Scientific, catalog number MA1-81357) for 1 hour and washed three times for 10 minutes in TBST solution. Protein bands were imaged using an Immuno-Star™ Reagents (Bio-Rad, Hercules, CA) and X-ray film.

### Allele Sequencing and Expression Analysis

DNA was isolated from ground frozen leaf tissue using the GeneJET Plant Genomic DNA Purification Kit (Thermo Scientific™). Primers were designed throughout the genes of interest and fragments were amplified from genomic DNA using Q5 2X Master Mix (NEB), then Sanger sequenced at the Cornell Biotechnology Resource Center. RNA was isolated from the primary leaf of a 10-day old plant using the RNeasy Plant Mini Kit (QIAGEN) after freezing and grinding. RNA samples were quantified using a NanoDrop™ spectrophotometer (Thermo Scientific™) and 500 ng of RNA from each sample were used to make cDNA with SuperScript III Reverse Transcriptase (Invitrogen) and oligo dT primers. DreamTaq™ DNA Polymerase (Thermo Scientific™) was used for 30-cycle PCRs of 1 μl of cDNA or 50 ng of gDNA template. Eight microliters of the PCR products were then visualized in a 1% agarose gel. Samples chosen for expression and sequence analysis were done so based on NAM population parent lines and to encompass two or more lines for all phenotypes.

### Electrolyte leakage assays in *N. benthamiana*

Electrolyte leakage assays were performed as described previously (Kim et al., 2016). In brief, after infiltration of *Agrobacterium* strains into *N. benthamiana*, leaf discs were collected from the infiltrated area using a cork borer (5 mm diameter) 2 h post dexamethasone application. Four leaf discs from four individual leaves of four different plants were included for each replication. The leaf discs were washed three times with distilled water and floated in 5 ml of distilled water supplemented with 0.001% Tween 20 (Sigma-Aldrich). Conductivity was monitored using a Traceable Pen Conductivity Meter (VWR) at the indicated time points after dexamethasone induction.

### Immunoprecipitation assay in *N. benthamiana*

Frozen *N. benthamiana* leaf tissue (four leaves) was ground in 1 ml of IP buffer (50 mM Tris-HCl [pH 7.5], 150 mM NaCl, 10% Glycerol, 1 mM DTT, 1 mM EDTA, 1% NP40, 0.1% Triton X-100, 1% plant protease inhibitor cocktail [Sigma-Aldrich], and 1% 2,2’-dipyridyl disulfide [Chem-Impex]) using a ceramic mortar and pestle and gently rotated for 1 hour at 4°C. The samples were centrifuged at 10,000 x g for 10 minutes at 4°C twice to remove plant debris. Five hundred microliters of the clarified extract were then incubated with 10 μl of GFP-Trap A (Chromotek) α-GFP bead slurry overnight at 4°C with constant end-over-end rotation. After overnight incubation, the α-GFP beads were pelleted by centrifugation at 4000 x g for 1 minute at 4°C and washed five times with 500 μl of IP wash buffer. Eighty microliters of the immunocomplexes were resuspended in 20 μl of 5X SDS loading buffer, and the mixture was boiled at 95°C for 10 minutes. All protein samples were resolved on a 4-20% gradient Precise™ Protein Gels (Thermo Scientific, Waltham, MA) and separated at 185V for 1 hour in 1X Tris/Glycine/SDS running buffer. Total proteins were transferred to a nitrocellulose membrane (GE Water and Process Technologies, Trevose, PA). Membranes were blocked with 5% Difco™ Skim Milk (BD, Franklin Lakes, NJ) overnight at 4°C. Proteins were detected with 1:5,000 horseradish peroxidase-conjugated anti-HA antibody (rat monoclonal, Roche, catalog number 12013819001) or 1:5,000 monoclonal mouse anti-GFP antibody (Novus Biologicals, Littleton, CO, catalog number NB600-597), washed in 1X Tris-buffered saline (TBS; 50 mM Tris-HCl, 150 mM NaCl, pH 7.5) solution containing 0.1% Tween 20 (TBST) overnight and incubated with 1:5,000 horseradish peroxidase-conjugated goat anti-mouse antibody (abcam, Cambridge, MA catalog number ab6789). The nitrocellulose membranes were washed three times for 15 minutes in TBST solution and protein bands were imaged using an Immuno-Star™ Reagents (Bio-Rad, Hercules, CA) or Supersignal^®^ West Femto Maximum Sensitivity Substrates (Thermo Scientific, Waltham, MA) and X-ray film.

### DAB assay for hydrogen peroxide accumulation in wheat

Hydrogen peroxide accumulation was detected following the protocol described by Liu et al. (2012) and Thordal-Christensen et al. (1997). In brief, 0.01 g of DAB powder (Sigma-Aldrich) was dissolved in 10 ml of distilled water (pH 3.6) and incubated at 37°C for 1 hour on a New Brunswick orbital shaker to dissolve the DAB powder. Wheat leaf segments were harvested from the infiltrated leaves 3 days post inoculation, (10 plants per treatment, experiment performed twice), immersed immediately in DAB solution and vacuum infiltrated for 10 seconds. The samples were wrapped in aluminum foil and incubated overnight in the dark. After overnight incubation, the stained leaf tissue was gently rinsed with distilled water, submerged in 70% ethanol and incubated at 70°C to clear the chlorophyll. The cleared leaves were rinsed and stored in a lactic acid/glycerol/H_2_O solution (1:1:1, v/v/v) for photography. Wheat leaves inoculated with 10 mM MgCl_2_ (mock) or *P. syringae* DC3000(D36E) expressing AvrPphB(C98S) were used as controls.

## Acknowledgements

The authors thank Alex Ollhoff, Ana M. Poets, Gary Muehlbauer and Kevin Smith at the University of Minnesota, St. Paul, MN for pre-publication access to Rasmusson spring NAM populations and associated SNP marker data and maps; Priyanka Tyagi and Gina Brown-Guedira, USDA-ARS, Raleigh, NC for generating GBS marker data; Alan Collmer and Hai-Lei Wei, Cornell University, for the use of *Pseudomonas syringae* DC3000(D36E); Greg Fuerst, USDA-ARS, Ames, IA, for generation, maintenance, and distribution of barley germplasm; Hana Zandkarimi and Leina Joseph, Indiana University, for technical assistance; and the USDA Wheat Germplasm Collection and Scot Hulbert, Washington State University, for wheat seed.

This research was supported in part by a National Science Foundation Plant Genome Research Program grant 13-39348 to RPW, AJB, and RWI, NSF Grant Number IOS-1551452 to RWI, and USDA-Agricultural Research Service project 3625-21000-060-00D to RPW. The funders had no role in study design, data collection and analysis, decision to publish, or preparation of the manuscript. Mention of trade names or commercial products in this publication is solely for the purpose of providing specific information and does not imply recommendation or endorsement by the U.S. Department of Agriculture or the National Science Foundation. USDA is an equal opportunity provider and employer.

